# Neutrophil-Derived S100A8/A9 Drives Inflammation that Promotes *Dnmt3a*-Mutant Hematopoiesis

**DOI:** 10.64898/2026.06.26.734778

**Authors:** Jayna J. Mistry, Natalie R. Fournier, Griffin J. Nye, Jennifer J. Trowbridge

**Affiliations:** The Jackson Laboratory, Bar Harbor, ME, USA

**Author notes:** Corresponding Author: Jennifer J. Trowbridge, The Jackson Laboratory, 600 Main Street, Bar Harbor, ME 04609. **Authorship** J.J.M. and J.J.T. designed experiments. J.J.M., N.R.F. and G.J.N. performed experiments, and J.J.M., N.R.F., G.J.N. and J.J.T. analyzed data. J.J.M. and J.J.T. interpreted data and wrote the manuscript. All authors edited the manuscript.

## Abstract

Clonal hematopoiesis (CH) is an age-associated condiion defined by over-representation of hematopoietic stem cells (HSCs) and their progeny carrying somatic mutations or variants that confer a selective advantage. CH is associated with increased risk of hematologic malignancies (1), cardiovascular disease and inflammatory bone loss (2, 3). Chronic inflammation is increasingly recognized as a central mediator of CH-mutant hematopoietic stem and progenitor cell (HSPC) expansion underlying CH (4). DNA methyltransferase 3a (*Dnmt3a*)-mutant cells produce higher levels of tumor necrosis factor-α (TNFα) and interleukin-6 (IL-6) (5), and blocking these pathways reduces the competitive advantage of *Dnmt3a*-mutant HSPCs (4, 6). The upstream mediators initiating inflammatory signaling in CH are unknown. Strong candidates are S100A8 and S100A9, members of the S100 calcium-binding protein family that regulate inflammatory signaling in the hematopoietic system. These proteins form a heterodimer complex and activate innate immune signaling through receptors including Toll-like receptor 4 (TLR4) and the receptor for advanced glycation end products (RAGE) (7). S100A8/A9 signaling promotes production of pro-inflammatory cytokines and inflammasome activation leading to poor prognosis in myelodysplastic syndrome and myeloproliferative neoplasms (8, 9). Across multiple myeloid malignancies, neutrophils are the primary bone marrow (BM) source of this alarmin (8, 10, 11) and pharmacologic inhibition of S100A9 with tasquinimod reduces disease severity without disrupting normal hematopoiesis (10, 12). Given the role of S100A8/A9 in establishing an inflammatory milieu, here we investigated the role of S100A8/A9 in *Dnmt3a*-mutant hematopoiesis. We identify neutrophils as a major source of elevated S100A8/A9 in the BM of *Dnmt3a*-mutant mice and this increase correlates with production of the inflammatory cytokines TNFα and IL-6. We show that tasquinimod reduces TNFα and IL-6 levels and selectively reduces the *Dnmt3a*-mutant HSPC compartment.

---

To characterize S100A8/A9 in the context of *Dnmt3a*-mutant hematopoiesis, we employed a transplantation model in which BM from *Dnmt3a*-mutant or Mx1-Cre control mice were transplanted into busulfan conditioned 2-4 month wild-type recipient animals (**Figure 1A**). After 12 weeks of engraftment, mice with *Dnmt3a*-mutant hematopoiesis exhibited elevated levels of the alarmins S100A8 and S100A9 in the extracellular BM fluid compared to controls (**Figure 1B**). To determine the putative cellular source of S100A8/A9, we analyzed published single-cell RNA sequencing (scRNA-seq) data from mice transplanted with *Dnmt3a*-mutant or Fgd5-CreERT2 control BM cells (5). We observed that *S100a8* and *S100a9* transcripts were most abundant within neutrophil populations, particularly mature neutrophils and immature granulocytes (**Supplementary Figure 1A**), consistent with neutrophils being the primary source of this alarmin in the BM (8, 10, 11). Both *S100a8* and *S100a9* were significantly increased in neutrophils and immature granulocytes in the *Dnmt3a*-mutant condition compared to their control counterparts (**Figure 1C**). We next assessed whether increased S100A8/A9 expression was also evident at the protein level under steady-state conditions. We utilized young adult (2-4mo) *Dnmt3a*-mutant or Mx1-Cre control mice (13). Intracellular flow cytometry revealed increased expression of S100A8/A9 in *Dnmt3a*-mutant neutrophils (CD11b^+^ Gr-1^hi^) and immature granulocytes (CD11b^+^ Gr-1^int^) compared to controls (**Figure 1D-E**, gating strategy shown in **Supplementary Figure 1B**) with no differences in the frequencies of these populations between *Dnmt3a*-mutant and control mice (**Supplementary Figure 1C**). These data show that the *Dnmt3a* mutation intrinsically increases expression of alarmins in neutrophil populations.

**Figure 1.**
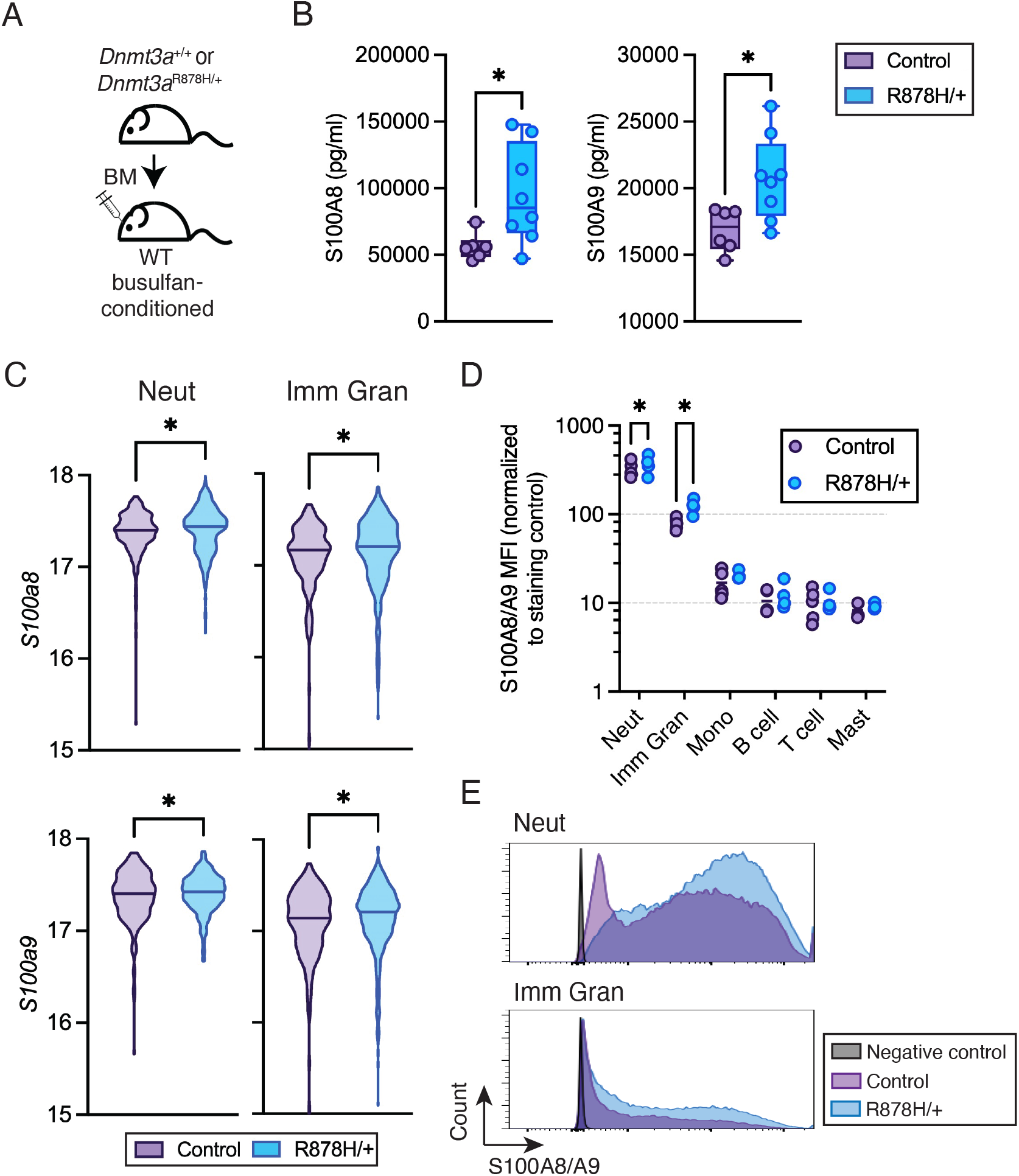
*Dnmt3a*-mutant hematopoiesis is associated with increased neutrophil expression of S100A8/A9. **(A)** Schematic of busulfan-conditioned transplant experimental design. **(B)** Concentration of S100A8 and S100A9 in BM extracellular fluid of control or *Dnmt3a*^R878H/+^ transplant mice. \**P* < 0.05 by Mann-Whitney test (n = 6 control, n = 8 *Dnmt3a*^R878H/+^ biological replicates). **(C)** Violin plots of single cell expression of *S100a8* (top) and *S100a9* (bottom) in neutrophils (left) and immature granulocytes (right) in control or *Dnmt3a*^R878H/+^ transplanted mice. Y-axis shows expression range within detected cells. \**P <* 0.05 by unpaired *t* test with Welch’s correction (cells isolated from n = 4 biological replicate mice per genotype). **(D**,**E)** Quantification (**D**) and representative histograms (**E**) of intracellular flow cytometric analysis of S100A8/A9 in BM populations in steady-state control and *Dnmt3a*^R878H/+^ mice. Neut; neutrophils (CD11b^+^ Gr1^hi^), Imm Gran; immature granulcoytes (CD11b^+^ Gr1^int^), Mono; monocytes (CD11b^+^ Gr1^lo^), B cells (B220^+^ CD11b^-^), T cells (CD3e^+^ B220^-^ CD11b^-^), Mast cells (FCeR1a^+^ B220^-^ CD11b^-^). \**P* < 0.05 by two-way ANOVA with Sidak’s multiple comparisons test (n = 5 control, n = 4 *Dnmt3a*^R878H/+^ biological replicate mice).

We established an *in vitro* culture system to investigate the consequences of S100A8/A9 production by *Dnmt3a*-mutant cells on inflammatory cytokine production. Hematopoietic progenitor cells (c-Kit^+^) were isolated from steady-state control or *Dnmt3a*-mutant Mx1-Cre mice and cultured in conditions enabling progenitor cell differentiation for 7 days (**Figure 2A**). These cultures contained cell surface marker-defined neutrophils (CD11b^+^ Gr-1^hi^) and immature granulocytes (CD11b^+^ Gr-1^int^) at similar proportions (**Figure 2B**). Analysis of the conditioned media (CM) from these cultures revealed increased levels of S100A8 and S100A9 produced by *Dnmt3a*-mutant cells (**Figure 2C**), demonstrating that production of alarmins by *Dnmt3a*-mutant cells can be modeled *in vitro*. S100A9 levels in the CM correlated significantly with the levels of TNFα (r = 0.9229, p = 0.0011), IL-6 (r = 0.9428, p = 0.0005) and IL-1α (r = 0.7068, p = 0.0500) (**Figure 2D**), showing that S100A8/A9 is positively correlated with inflammatory cytokine production. To evaluate if S100A8/A9 drives elevated inflammatory cytokine production, we cultured hematopoietic progenitor cells (c-Kit^+^) from control or *Dnmt3a*-mutant mice in the presence or absence of the S100A9 inhibitor tasquinimod (**Figure 2E**). Tasquinimod significantly reduced TNFα and IL-6 production by *Dnmt3a*-mutant cells but did not alter IL-1α (**Figure 2F**). The lack of effect on IL-1α, which is released through cell damage-dependent mechanisms and not a canonical target of S100A9, supports that tasquinimod acts selectively to dampen the *Dnmt3a*-mutant S100A9-driven inflammatory program rather than suppressing cytokine production broadly.

**Figure 2.**
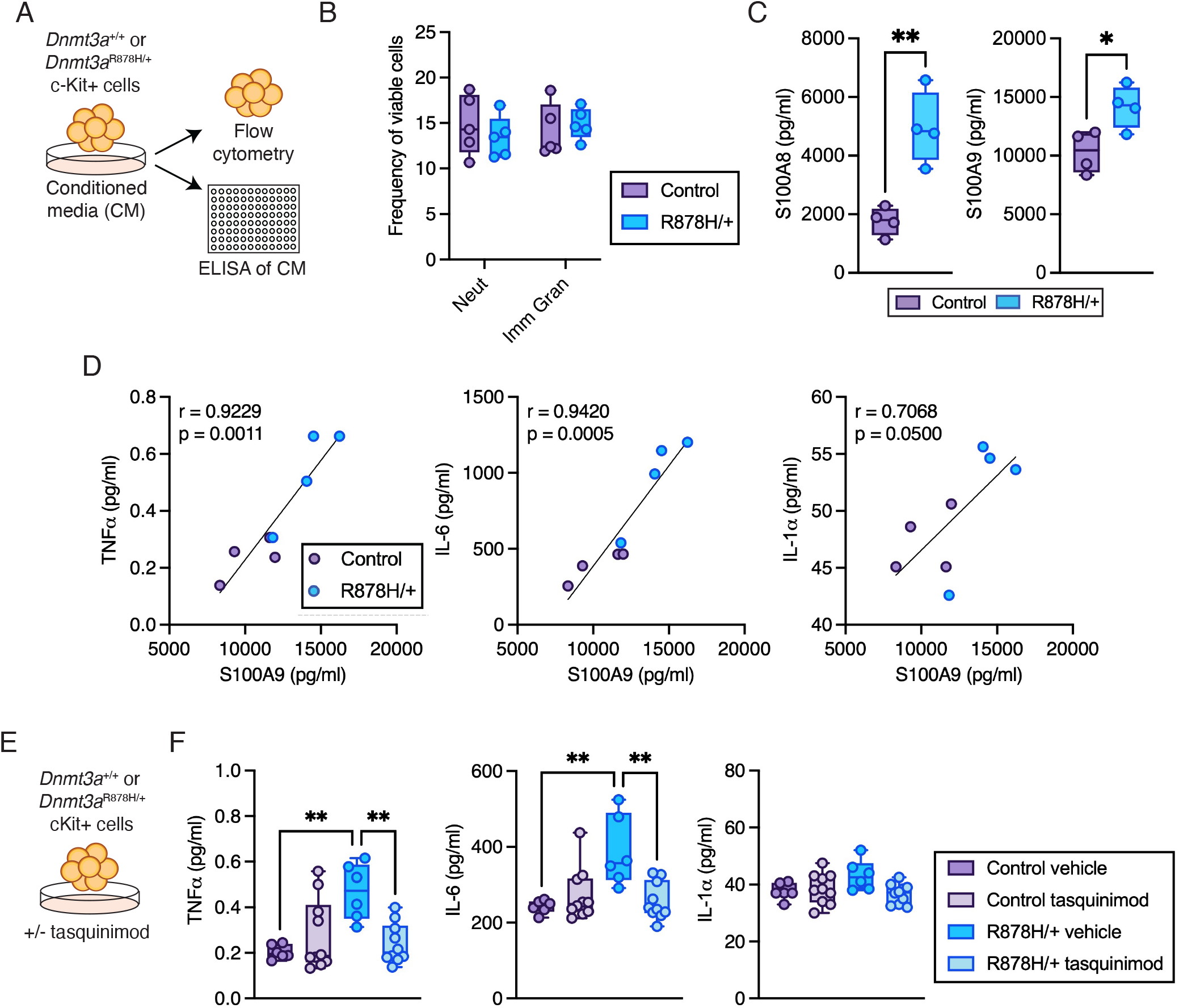
S100A8/A9 promotes inflammatory cytokine production *ex vivo*. **(A)** Schematic of conditioned media (CM) experimental design. **(B)** Proportion of neutrophils (CD11b^+^ Gr-1^hi^) and immature granulocytes (CD11b^+^ Gr-1^int^) after 7-day culture of c-Kit^+^ hematopoietic progenitors from control or *Dnmt3a*^R878H/+^ mice. Not significant by two-way ANOVA with uncorrected Fisher’s LSD (n = 4 biological replicate mice per group). **(C)** Concentration of S100A8 and S100A9 in CM collected from 7-day cultures. \**P* < 0.05, ** *P*< 0.01 by unpaired *t* tests (n = 4 biological replicate mice per group). **(D)** Correlation between concentration of TNFα (left), IL-6 (center) or IL-1α (right) with the concentration of S100A9 in CM collected from 7-day cultures. Significance values calculated by two-tailed Pearson correlation (n = 4 biological replicate mice per group). **(E)** Schematic of culture experiment in which control or *Dnmt3a*^R878H/+^ c-Kit^+^ hematopoietic cells were cultured for 7 days in the presence or absence of tasquinimod followed by collection of CM. **(F)** Concentration of TNFα (left), IL-6 (center) or IL-1α (right) in CM. \*\**P* <0.01 by Kruskal-Wallis test with uncorrected Dunn’s test (n = 6 control vehicle, n = 10 control tasquinimod, n = 6 *Dnmt3a*^R878H/+^ vehicle, n = 10 *Dnmt3a*^R878H/+^ tasquinimod).

We next examined whether inhibition of S100A8/A9-driven inflammatory cytokine production impacts the competitive advantage of *Dnmt3a-*mutant cells. We first used a competitive *ex vivo* co-culture assay, whereby CD45.2^+^ control or *Dnmt3a*-mutant c-Kit^+^ progenitors were cultured with CD45.1^+^ wild-type competitor c-Kit^+^ progenitor cells in the presence or absence of tasquinimod for 7 days (**Figure 3A**, gating strategy shown in **Supplementary Figure 2A**). *Dnmt3a*-mutant hematopoietic cells had a competitive advantage over their wild-type counterparts, resulting in an increased proportion of CD45.2^+^ cells compared to control (**Figure 3B**). This overall competitive advantage was not significantly altered by tasquinimod treatment, and no differences in proportion of neutrophils or immature granulocytes (**Supplementary Figure 2B**) were observed. In contrast, the proportion of *Dnmt3a*-mutant HSPCs (Lin^−^ Sca-1^+^ c-Kit^+^) was increased relative to control and significantly reduced by tasquinimod (**Figure 3C**). To test whether S100A8/A9 augments *Dnmt3a*-mutant hematopoiesis *in vivo*, we treated competitive transplant recipients with tasquinimod continuously in their drinking water for 2 weeks (**Figure 3D**, gating strategy shown in **Supplementary Figure 2C**). Tasquinimod reduced the frequency of macrophages (CD11b^+^ Ly6G^-^ Ly6C^-^ F4/80^+^) and monocytes (CD11b^+^ Ly6G^-^ Ly6C^-^ F4/80^-^) in the peripheral blood (PB) of both control and *Dnmt3a*-mutant recipients (**Supplementary Figure 2D**), consistent with the myeloid modulatory activity of tasquinimod (14) and confirming that the drug was biologically active *in vivo*. In the BM, we found that *Dnmt3a*-mutant hematopoietic cells gave rise to higher competitive chimerism in recipient mice compared to control and tasquinimod treatment significantly reduced the proportion of *Dnmt3a*-mutant cells without affecting control cells (**Figure 3E**, left panel). Furthermore, *Dnmt3a*-mutant cells in the BM contained a higher proportion of phenotypic HSCs (Lin^-^ Sca-1^+^ c-Kit^+^ Flt3^-^ CD150^+^ CD48^-^) and this was significantly reduced with tasquinimod (**Figure 3E**, right panel). Prolonged treatment with tasquinimod was limited by dermatitis, a spontaneous condition of the C57BL/6 mouse strain background that occurred across groups but was more severe in tasquinimod-treated mice of both genotypes. A small cohort of surviving mice evaluated after 10 weeks of tasquinimod treatment showed similar directional changes. In the BM, at the bulk cell level and at the phenotypic HSC level, *Dnmt3a*-mutant hematopoietic cells had higher chimerism and tasquinimod-treated mice showed a trend toward reduction (**Figure 3F**). Together, these data indicate that S100A8/A9 supports the short-term competitive advantage of *Dnmt3a*-mutant HSPC in the BM.

**Figure 3.**
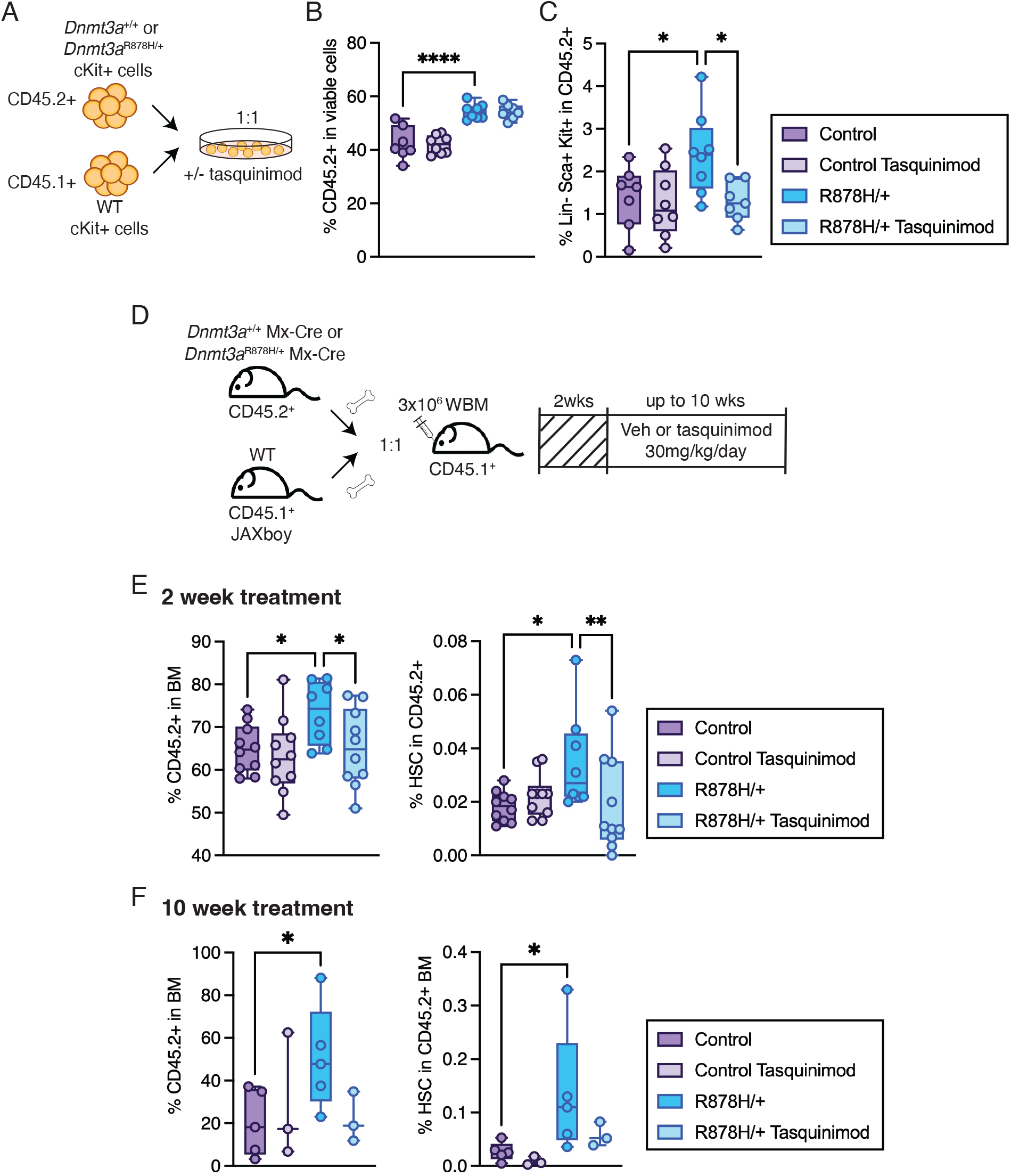
Inhibition of S100A8/A9 signaling reduces *Dnmt3a*-mutant HSPC expansion. **(A)** Schematic of competitive co-culture assay in which CD45.2^+^ control or *Dnmt3a*^R878H/+^ c-Kit^+^ hematopoietic cells were cultured with CD45.1^+^ competitor cells in the presence or absence of tasquinimod for 7 days. **(B)** Frequency of CD45.2^+^ cells after competitive 7-day co-culture with or without tasquinimod. **(C)** Frequency of phenotypic HSPCs (Lin^-^ Sca-1^+^ c-Kit^+^) in CD45.2^+^ after competitive 7-day co-culture with or without tasquinimod. \**P* < 0.05, \*\*\*\**P* < 0.0001 by Brown-Forsythe and Welch’s ANOVA with unpaired t test (n = 7 control vehicle, n = 8 control tasquinimod, n = 8 *Dnmt3a*^R878H/+^ vehicle, n = 7 *Dnmt3a*^R878H/+^ tasquinimod). **(D)** Schematic of the competitive transplantation experiment with tasquinimod provided ad libitum in the drinking water. **(E)** Frequency of donor (CD45.2^+^) cells (left) and donor-derived HSCs (Lin^-^ Sca-1^+^ c-Kit^+^ Flt3^-^ CD150^+^ CD48^-^) (right) in the BM of recipient mice receiving control or tasquinimod-supplemented water for 2 weeks. \**P* < 0.05, \*\**P* < 0.01 by Kruskal-Wallis test with uncorrected Dunn’s test (n = 10 control vehicle, n = 10 control tasquinimod, n = 8 *Dnmt3a*^R878H/+^ vehicle, n = 10 *Dnmt3a*^R878H/+^ tasquinimod). **(F)** Frequency of donor (CD45.2^+^) cells (left) and donor-derived HSCs (Lin^-^ Sca-1^+^ c-Kit^+^ Flt3^-^ CD150^+^ CD48^-^) in the BM of recipient mice receiving control or tasquinimod-supplemented water for 10 weeks. \**P* < 0.05 by Kruskal-Wallis test with uncorrected Dunn’s test (n = 5 control vehicle, n = 3 control tasquinimod, n = 5 *Dnmt3a*^R878H/+^ vehicle, n = 3 *Dnmt3a*^R878H/+^ tasquinimod).

These observations build upon and extend a growing body of evidence demonstrating that CH is characterized by a pro-inflammatory BM microenvironment. Our data identify neutrophil-derived S100A8/A9 as a driver of inflammation and demonstrate that pharmacologic inhibition of this axis reduces both inflammatory cytokine production and the *Dnmt3a*-mutant HSPC compartment. Notably, targeting the alarmin axis is advancing toward clinical translation in related myeloid diseases. Tasquinimod is currently being evaluated in a Phase I/II trial in myelofibrosis (HOVON 172, NCT06605586) (12) and preclinical evidence supports its efficacy in MDS and AML models as well (11, 15). Our data provide a pre-malignant rationale for this approach, suggesting that intervening at the CH stage to disrupt S100A8/A9-driven inflammatory signaling may suppress mutant HSPC expansion before overt disease develops. Future investigations should also explore whether S100A8/A9 released by *Dnmt3a*-mutant neutrophils also remodels the BM stromal niche, as has been demonstrated in MDS and myelofibrosis (11, 12).

## Figure Legends

**Supplementary Figure 1. (A)** Single-cell RNA-seq (scRNA-seq) analysis of *S100a8* and *S100a9* across hematopoietic and non-hematopoietic cell populations derived from busulfan-conditioned transplant (n = 4 individual biological replicate mice per genotype). Circle size represents frequency of population in which transcript was detected, color intensity represents scaled mean expression. HSC/MPP: hematopoietic stem and multipotent progenitor cells, MPP-GM: granulocyte-macrophage-primed multipotent progenitor cells, MPP-Ly: lymphoid-primed multipotent progenitor cells, CycProg: cycling progenitor cells, GMP: granulocyte-macrophage progenitor cells, MEP: megakaryocyte-erythroid progenitor cells, EryPro: erythroid progenitor cells, Ery: erythroblasts, MEoBasoPro: mast-eosinophil-basophil progenitor cells, MonoPro: monocyte progenitor cells, Mono/DC: monocytes and dendritic cells, Mono: monocytes, Mac: macrophages, DC: dendritic cells, NeutPro: neutrophil progenitor cells, ImmGran: immature granulocytes, Neut: neutrophils, NK/T: natural killer and T cells, PreB: pre-B cells, ProB: pro-B cells, B: B cells, EC-Sin: sinusoidal endothelial cells, EC-Art: arteriolar endothelial cells, OB: osteoblasts, MSC: mesenchymal stromal cells. **(B)** Representative gating strategy for fixed BM cells to evaluate intracellular S100A8/A9 staining. **(C)** Flow cytometric quantification of neutrophil (CD11b^+^ Gr1^hi^) and immature granulocyte (CD11b^+^ Gr1^int^) frequencies in the BM of steady-state control or *Dnmt3a*^R878H/+^ mice. Not significant by two-way ANOVA with uncorrected Fisher’s LSD (n = 6 biological replicate mice per genotype).

**Supplementary Figure 2. (A)** Representative gating strategy in the analysis of cells after competitive 7-day co-culture. **(B)** Frequency of cell surface marker-defined neutrophils (CD11b^+^ Gr-1^hi^) and immature granulocytes (CD11b^+^ Gr-1^int^) in CD45.2^+^ cells after competitive 7-day co-culture with or without tasquinimod. Not significant by by Brown-Forsythe and Welch’s ANOVA with unpaired t test (n = 7 control vehicle, n = 8 control tasquinimod, n = 8 *Dnmt3a*^R878H/+^ vehicle, n = 7 *Dnmt3a*^R878H/+^ tasquinimod). **(C)** Representative gating strategy for BM analysis of mice receiving control or tasquinimod-supplemented water. **(D)** Frequency of macrophages (CD11b^+^ Ly6G^-^ Ly6C^-^ F4/80^+^) (left) and monocytes (CD11b^+^ Ly6G^-^ Ly6C^-^ F4/80^-^) (right) in the PB of mice receiving control or tasquinimod-supplemented water for 2 weeks. \**P* < 0.05, \*\**P* < 0.01, \*\*\**P* <0.001 by Kruskal-Wallis test with uncorrected Dunn’s test (n = 7 control vehicle, n = 5 control tasquinimod, n = 7 *Dnmt3a*^R878H/+^ vehicle, n = 5 *Dnmt3a*^R878H/+^ tasquinimod).

## Methods

### Mice

All animal work was approved by the Institutional Animal Care and Use Committee at The Jackson Laboratory (Bar Harbor, ME) (protocol #12004-01). The strains C57BL/6J (RRID:IMSR_JAX:000664), B6.SJL-*Ptprc*a (RRID:IMSR_JAX:002014, referred to as CD45.1^+^), and C57BL/6J-*Ptprc*em6Lutzy (RRID:IMSR_JAX:033076, referred to as CD45.1^+^ JAXboy) were obtained from and housed within The Jackson Laboratory. B6(Cg)-*Dnmt3a*tm1Trow/J (RRID:IMSR_JAX:032289, referred to as *Dnmt3a*^R878H/+^ or *Dnmt3a*-mutant) mice were crossed to B6.Cg-Tg(Mx-cre)1Cgn/J (RRID:IMSR_JAX:003556, referred to as Mx1-Cre) mice or to the combination of C57BL/6N-*Fgd5*tm3(cre/ERT2)Djr/J (RRID:IMSR_JAX:027789, referred to as Fgd5-CreERT2) and B6.Cg-*Gt(ROSA)26Sor*tm14(CAG-tdTomato)Hze/J (RRID:IMSR_JAX:007914) mice. Mice carrying the Mx-Cre allele were given 15 mg/kg polyinosinic-polycytidylic acid (poly(I:C), InvivoGen) by intraperitoneal (IP) injection every other day for a total of five doses between 2–4 months of age. Mice carrying the Fgd5-CreER allele were given 125 mg/kg/day tamoxifen (TAM) by oral gavage for 3 days. Following poly(I:C) or TAM treatment, mice recovered for >5 weeks prior to use in experiments. Mice were housed in cages under specific pathogen-free conditions in a 12/12-hour light/dark, temperatures of 65–75°F with 40– 60% humidity, food and water provided *ad libitum*. All mice were used at 2-4 months of age, and both male and female mice were used except for transplantation experiments where female mice were used.

### BM transplantation

For busulfan-conditioned transplant experiments, 3×10^6^ BM cells from *Dnmt3a*^R878H/+^ Mx1-Cre, *Dnmt3a*^R878H/+^ Fgd5-CreERT2 LSL-tdTomato, or respective control donor mice were transplanted by retro-orbital (RO) injection into wild-type 9 month old CD45.1^+^ recipient mice conditioned with busulfan (20 mg/kg/day IP for 3 days) prior to transplant. For competitive transplantation experiments, CD45.2^+^ donor BM cells were mixed 1:1 with CD45.1^+^ JAXboy or CD45.1^+^ competitor BM cells and transplanted into 3 month CD45.1^+^ recipient mice conditioned using 1100 rads of X-ray irradiation (split dose, 3-4 hours apart). PB chimerism was monitored every 4 weeks after transplant by flow cytometry of PB collected by retro-orbital or submandibular blood collection. In some experiments, tasquinimod (Biorbyt, Cat. orb1300548) was administered to recipient mice via the drinking water at 30 mg/kg/day. Tasquinimod was reconstituted in DMSO (Sigma-Aldrich, Cat. D2650) and dissolved in 3% sucrose (Sigma-Aldrich, Cat. S7903-1KG) and 2% PEG300 (MedChemExpress, Cat. HY-Y0873). Vehicle-treated groups received DMSO, PEG300-treated water with 3% sucrose. Treatment started 2 weeks after transplantation and continued for up to 10 weeks. In some cases, treatment duration was limited by dermatitis that was more severe with tasquinimod treatment, necessitating early sacrifice of mouse cohorts.

### Luminex

For multiplexed ELISA (Luminex) experiments following transplant with *Dnmt3a*^R878H/+^ Mx-Cre BM, one femur and one tibia were cut in half then spun at 4000xg for 3 seconds into 100 μL of PBS. Analyte levels were measured using custom Luminex assays (R&D Systems) for mouse S100A8, S100A9, TNFα, IL-6, and IL-1α.

### Flow cytometry of BM and PB

Single-cell suspensions of BM were prepared by filtering crushed femurs, tibias, and iliac crests from each mouse. PB was collected as described above. Red cells were lysed from BM and PB samples using 1X red blood cell lysis buffer (ThermoFisher). Cells were stained with a combination of fluorochrome-conjugated antibodies: CD45.1 BV650 (clone A20, BioLegend Cat. 110736, 1:200), CD45.2 A700 (clone 104, BioLegend Cat. 109822, 1:50), c-Kit BV650 (clone 2B8, BD Biosciences Cat. 563399, 1:200) or c-Kit APC (clone 2B8, BD Biosciences Cat. 553356, 1:200), Sca-1 BV510 (clone D7, BioLegend Cat. 108129, 1:200), CD150 BV786 (clone TC15-12F12.2, BioLegend Cat. 115937, 1:200), CD48 PE-Cy7 (clone HM48-1, BioLegend Cat. 103424, 1:200), Flt3/CD135 BV421 (clone A2F10, BioLegend Cat. 135313, 1:50), CD34 PE-Cy5 (clone MEC14.7, BioLegend Cat. 119311, 1:50) or CD34 FITC (clone RAM34, BD Biosciences Cat. 553733, 1:50), FcgR BV711 (clone 93, BioLegend Cat. 101337, 1:200), CD11b BV605 (clone M1/70, BioLegend Cat. 101257, 1:200), F4/80 PE (clone BM8, BioLegend Cat. 123109, 1:200), B220 BUV496 (clone RA3-6B2, BD Biosciences Cat. 612950, 1:200), FceRIa PE (clone MAR-1, BioLegend Cat. 134307, 1:200), CD3e PerCP-Cy5.5 (clone 145-2C11, BioLegend Cat. 100328, 1:200), Gr-1 APC-Cy7 (clone RB6-8C5, BioLegend Cat. 108423, 1:200), Ly6G APC (clone 1A8, BioLegend, Cat. 127613, 1:200), Ly6C BV605 (clone HK1.4, BioLegend Cat. 128035, 1:200), a mature lineage (Lin) marker mix including B220 APC (clone RA3-6B2, BioLegend Cat. 103211, 1:200) or B220 PE-Cy5 (clone RA3-6B2, BioLegend Cat. 103210, 1:200), CD4 APC (clone RM4-5, BioLegend Cat. 100515, 1:200), or CD4 PE-Cy5 (clone RM4-5, BioLegend Cat. 100514, 1:200), CD5 APC (clone 53-7.3, BioLegend Cat. 100625, 1:200) or CD5 PE-Cy5 (clone 53-7.3, BioLegend Cat. 100610, 1:200), CD8a APC (clone 53-6.7, BioLegend Cat. 100711, 1:200) or CD8a PE-Cy5 (clone 53-6.7, BioLegend Cat. 100710, 1:200) and Ter-119 APC (clone Ter119, BioLegend Cat. 116211, 1:200) or Ter-119 PE-Cy5 (clone Ter119, BioLegend Cat. 116210, 1:200), and a viability stain (DAPI). For intracellular S100A8/A9 staining, cells were fixed and permeabilized using the FIX & PERM Cell Permeabilization Kit (ThermoFisher) per the manufacturer’s instructions prior to intracellular antibody staining with rabbit anti-mouse S100A8/A9 (clone RM1038, Abcam Cat. ab288715, 1:500) and goat anti-rabbit AlexaFluor 488 (Abcam Cat. ab150081, 1:5000). S100A8/A9 MFI values were normalized to the mean MFI of negative staining controls collected within the same experimental batch, prior to statistical comparison. Antibody cocktails were incubated with BM cells for ≥30 min at 4°C. Stained cells were analyzed on a FACSymphony A5 (BD). FlowJo V10 was used for data analysis.

### Hematopoietic progenitor cell culture

BM was prepared by isolating and filtering crushed pooled tibias, femurs, and iliac crests from each mouse. HSPCs (c-Kit^+^) were isolated using CD117 MicroBeads (Miltenyi) and cultured in StemSpan SFEM II (StemCell Technologies) supplemented with 10 ng/mL recombinant murine SCF (StemCell Technologies) for 7 days at 37°C in 5% CO_2_. Cells were harvested for analysis by flow cytometry as described above. To retain CM for analysis, following culture the cells were centrifuged at 300 × g for 5 min at 4°C, supernatant collected and re-centrifuged at 1,300 × g for 10 min at 4°C. This CM was used for Luminex assays described above. For competitive co-culture assays, CD45.2^+^ c-Kit^+^ cells from control or *Dnmt3a*-mutant mice were co-cultured with CD45.1^+^ JAXboy competitor c-Kit^+^ cells at a 1:1 ratio (30000 CD45.2^+^ c-Kit^+^ cells with CD45.1^+^ JAXboy competitor c-Kit^+^ cells) for 7 days prior to assessment by flow cytometry as described above. In some experiments, competitive co-cultures were treated with either 50 μM tasquinimod (Biorbyt, Cat. orb1300548) or DMSO (Sigma-Aldrich, Cat. D2650) vehicle control at cell seeding and spiked in every 24□hours for 7 days. Cells were analyzed by flow cytometry as described above.

### Statistics

No statistical method was used to predetermine sample size. The investigators were not blinded to allocation during experiments and outcome assessment. All statistical tests were performed using Prism 11 (GraphPad) as described in figure legends. Differences among group means were considered statistically significant when *P* < 0.05. Sample size (n) represents the number of biological replicates.

## Acknowledgements

This work was supported by NIH grants R01DK118072 and R01AG069010, and an EvansMDS Discovery Research Grant to J.J.T, and U01AG077925 to J.J.T. This work was supported in part by the NIH/NCI Cancer Center Support Grant P30CA034196. J.J.T. was supported by a Leukemia & Lymphoma Society Scholar Award and The Dattels Family Endowed Chair. J.J.M. was supported by a Leukemia & Lymphoma Society Career Development Program Fellow Award and The Jackson Laboratory Scholar Award. We thank all members of the Trowbridge lab for experimental support. We thank The Jackson Laboratory’s Single Cell Biology, Genome Technologies, Clinical Chemistry and Flow Cytometry Scientific Services. We thank R. Schneider, H. Gleitz, A. Verma, R. Zhao, K. Young and M. Telpoukhovskaia for their input into this work.

## Data Availability

Published single-cell RNA sequencing data from mice transplanted with *Dnmt3a*-mutant or Fgd5-CreERT2 control BM cells(5) are available in the Gene Expression Omnibus under accession code GSE240686.

## References

1. Jaiswal S, Fontanillas P, Flannick J, Manning A, Grauman PV, Mar BG, Lindsley RC, Mermel CH, Burtt N, Chavez A, Higgins JM, Moltchanov V, Kuo FC, Kluk MJ, Henderson B, Kinnunen L, Koistinen HA, Ladenvall C, Getz G, Correa A, Banahan BF, Gabriel S, Kathiresan S, Stringham HM, McCarthy MI, Boehnke M, Tuomilehto J, Haiman C, Groop L, Atzmon G, Wilson JG, Neuberg D, Altshuler D, Ebert BL. Age-related clonal hematopoiesis associated with adverse outcomes. N Engl J Med. 2014 Dec 25;371(26):2488–98. PMID 25426837.

2. Jaiswal S, Natarajan P, Silver AJ, Gibson CJ, Bick AG, Shvartz E, McConkey M, Gupta N, Gabriel S, Ardissino D, Baber U, Mehran R, Fuster V, Danesh J, Frossard P, Saleheen D, Melander O, Sukhova GK, Neuberg D, Libby P, Kathiresan S, Ebert BL. Clonal Hematopoiesis and Risk of Atherosclerotic Cardiovascular Disease. N Engl J Med. 2017 Jul 13;377(2):111–121. PMID 28636844.

3. Wang H, Divaris K, Pan B, Li X, Lim JH, Saha G, Barovic M, Giannakou D, Korostoff JM, Bing Y, Sen S, Moss K, Wu D, Beck JD, Ballantyne CM, Natarajan P, North KE, Netea MG, Chavakis T, Hajishengallis G. Clonal hematopoiesis driven by mutated DNMT3A promotes inflammatory bone loss. Cell. 2024 Jul 11;187(14):3690–3711 e19. PMID 38838669.

4. Zioni N, Bercovich AA, Chapal-Ilani N, Bacharach T, Rappoport N, Solomon A, Avraham R, Kopitman E, Porat Z, Sacma M, Hartmut G, Scheller M, Muller-Tidow C, Lipka D, Shlush E, Minden M, Kaushansky N, Shlush LI. Inflammatory signals from fatty bone marrow support DNMT3A driven clonal hematopoiesis. Nat Commun. 2023 Apr 12;14(1):2070. PMID 37045808.

5. Mistry JJ, Young KA, Colom Diaz PA, Maestre IF, Levine RL, Trowbridge JJ. Mesenchymal Stromal Cell Senescence Induced by Dnmt3a -Mutant Hematopoietic Cells is a Targetable Mechanism Driving Clonal Hematopoiesis and Initiation of Hematologic Malignancy. bioRxiv. 2024 Mar 30. Epub 20240330. doi:10.1101/2024.03.28.587254. Cited in: Pubmed; PMID 38585779.

6. SanMiguel JM, Eudy E, Loberg MA, Young KA, Mistry JJ, Mujica KD, Schwartz LS, Stearns TM, Challen GA, Trowbridge JJ. Distinct Tumor Necrosis Factor Alpha Receptors Dictate Stem Cell Fitness versus Lineage Output in Dnmt3a-Mutant Clonal Hematopoiesis. Cancer Discov. 2022 Dec 2;12(12):2763–2773. PMID 36169447.

7. Chen B, Miller AL, Rebelatto M, Brewah Y, Rowe DC, Clarke L, Czapiga M, Rosenthal K, Imamichi T, Chen Y, Chang CS, Chowdhury PS, Naiman B, Wang Y, Yang D, Humbles AA, Herbst R, Sims GP. S100A9 induced inflammatory responses are mediated by distinct damage associated molecular patterns (DAMP) receptors in vitro and in vivo. PLoS One. 2015;10(2):e0115828. PMID 25706559.

8. Sallman DA, List A. The central role of inflammatory signaling in the pathogenesis of myelodysplastic syndromes. Blood. 2019 Mar 7;133(10):1039–1048. PMID 30670444.

9. Wang YH, Chen YJ, Lai YH, Wang MC, Chen YY, Wu YY, Yang YR, Tsou HY, Li CP, Hsu CC, Huang CE, Chen CC. Mutation-Driven S100A8 Overexpression Confers Aberrant Phenotypes in Type 1 CALR-Mutated MPN. Int J Mol Sci. 2023 May 14;24(10). PMID 37240094.

10. Leimkuhler NB, Gleitz HFE, Ronghui L, Snoeren IAM, Fuchs SNR, Nagai JS, Banjanin B, Lam KH, Vogl T, Kuppe C, Stalmann USA, Busche G, Kreipe H, Gutgemann I, Krebs P, Banz Y, Boor P, Tai EW, Brummendorf TH, Koschmieder S, Crysandt M, Bindels E, Kramann R, Costa IG, Schneider RK. Heterogeneous bone-marrow stromal progenitors drive myelofibrosis via a druggable alarmin axis. Cell Stem Cell. 2021 Apr 1;28(4):637–652 e8. PMID 33301706.

11. Wobus M, Weidner H, Wehner R, Baumann AL, Mobus K, Balaian E, Torngren M, Vahtola E, Eriksson H, Winter S, Platzbecker U, Chavakis T, Hofbauer LC, Rauner M, Bornhauser M, Sockel K. Preclinical efficacy of tasquinimod in myelodysplastic neoplasms: Restoring erythropoiesis and mitigating bone loss. Hemasphere. 2026 Apr;10(4):e70352. PMID 41987903.

12. Gleitz HFE, Fuchs SNR, Snoeren IAM, Boys C, Nagai J, Tejeda-Mora H, Kloker V, Pritchard JE, Bakker IJ, Gargallo Garasa M, Bindels E, Saez-Rodriguez J, Vogl T, Kramann R, Dugourd A, Costa IG, Schneider RK. Inhibiting the alarmin-driven hematopoiesis-stromal cell crosstalk in primary myelofibrosis ameliorates bone marrow fibrosis. Hemasphere. 2025 Aug;9(8):e70179. PMID 40823315.

13. Loberg MA, Bell RK, Goodwin LO, Eudy E, Miles LA, SanMiguel JM, Young K, Bergstrom DE, Levine RL, Schneider RK, Trowbridge JJ. Sequentially inducible mouse models reveal that Npm1 mutation causes malignant transformation of Dnmt3a-mutant clonal hematopoiesis. Leukemia. 2019 Jul;33(7):1635–1649. PMID 30692594.

14. Olsson A, Nakhle J, Sundstedt A, Plas P, Bauchet AL, Pierron V, Bruetschy L, Deronic A, Torngren M, Liberg D, Schmidlin F, Leanderson T. Tasquinimod triggers an early change in the polarization of tumor associated macrophages in the tumor microenvironment. J Immunother Cancer. 2015;3:53. Epub 20151215. PMID 26673090.

15. Cuminetti V, Boet E, Heugel M, Konieczny J, Bernal A, Gomez MJ, Grimolizzi F, Vilaplana-Lopera N, Ferre M, Villatoro A, Pandey DP, Torroja C, Taman H, Paulssen RH, Vogl T, Heckman CA, Vik A, Giovinazzo G, van Gastel N, Garcia P, Sanchez-Cabo F, Sarry JE, Arranz L. Succinate receptor 1 restricts hematopoiesis and prevents acute myeloid leukemia progression. Nat Commun. 2026 Feb 5;17(1). PMID 41644523.

