## Supplemental Figures for "Neutrophil-Derived S100A8/A9 Drives Inflammation that Promotes *Dnmt3a*-Mutant Hematopoiesis"

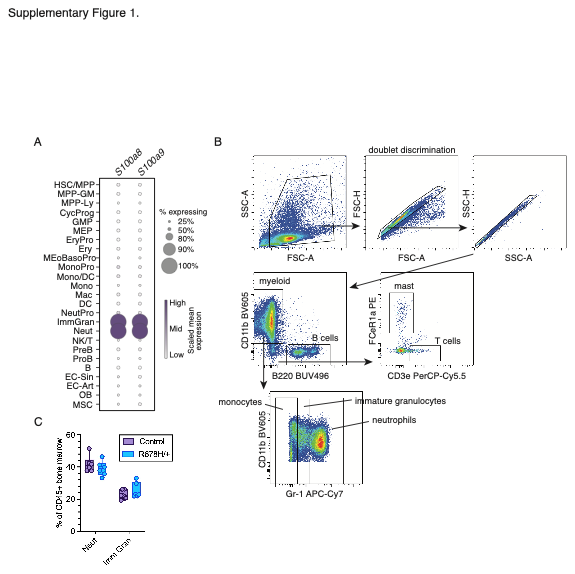

**Supplementary Figure 1. (A)** Single-cell RNA-seq (scRNA-seq) analysis of *S100a8* and *S100a9* across hematopoietic and non-hematopoietic cell populations derived from busulfan-conditioned transplant (n = 4 individual biological replicate mice per genotype). Circle size represents frequency of population in which transcript was detected, color intensity represents scaled mean expression. HSC/MPP: hematopoietic stem and multipotent progenitor cells, MPP-GM: granulocyte-macrophage-primed multipotent progenitor cells, MPP-Ly: lymphoid-primed multipotent progenitor cells, CycProg: cycling progenitor cells, GMP: granulocyte-macrophage progenitor cells, MEP: megakaryocyte-erythroid progenitor cells, EryPro: erythroid progenitor cells, Ery: erythroblasts, MEoBasoPro: mast-eosinophil-basophil progenitor cells, MonoPro: monocyte progenitor cells, Mono/DC: monocytes and dendritic cells, Mono: monocytes, Mac: macrophages, DC: dendritic cells, NeutPro: neutrophil progenitor cells, ImmGran: immature granulocytes, Neut: neutrophils, NK/T: natural killer and T cells, PreB: pre-B cells, ProB: pro-B cells, B: B cells, EC-Sin: sinusoidal endothelial cells, EC-Art: arteriolar endothelial cells, OB: osteoblasts, MSC: mesenchymal stromal cells. **(B)** Representative gating strategy for fixed BM cells to evaluate intracellular S100A8/A9 staining. **(C)** Flow cytometric quantification of neutrophil (CD11b^+^ Gr1^hi^) and immature granulocyte (CD11b^+^ Gr1^int^) frequencies in the BM of steady-state control or *Dnmt3a*^R878H/+^ mice. Not significant by two-way ANOVA with uncorrected Fisher’s LSD (n = 6 biological replicate mice per genotype).

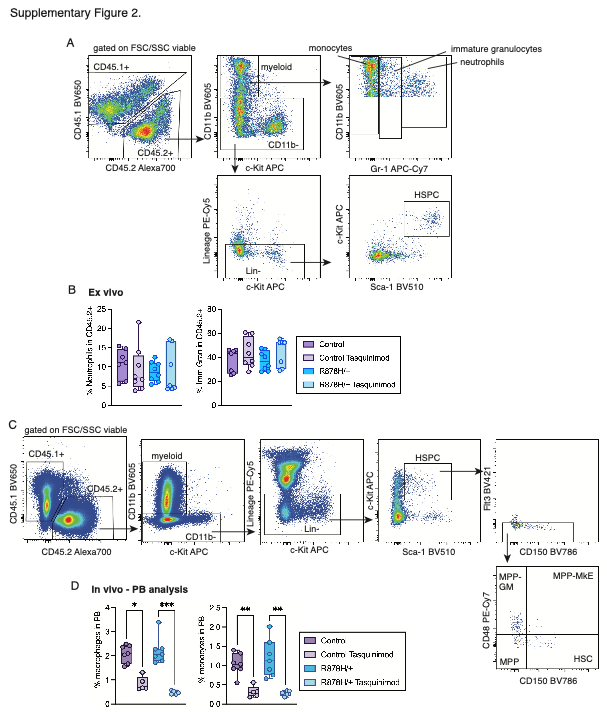

**Supplementary Figure 2. (A)** Representative gating strategy in the analysis of cells after competitive 7-day co-culture. **(B)** Frequency of cell surface marker-defined neutrophils (CD11b^+^ Gr-1^hi^) and immature granulocytes (CD11b^+^ Gr-1^int^) in CD45.2^+^ cells after competitive 7-day co-culture with or without tasquinimod. Not significant by by Brown-Forsythe and Welch’s ANOVA with unpaired t test (n = 7 control vehicle, n = 8 control tasquinimod, n = 8 *Dnmt3a*^R878H/+^ vehicle, n = 7 *Dnmt3a*^R878H/+^ tasquinimod). **(C)** Representative gating strategy for BM analysis of mice receiving control or tasquinimod-supplemented water. **(D)** Frequency of macrophages (CD11b^+^ Ly6G^-^ Ly6C^-^ F4/80^+^) (left) and monocytes (CD11b^+^ Ly6G^-^ Ly6C^-^ F4/80^-^) (right) in the PB of mice receiving control or tasquinimod-supplemented water for 2 weeks. **P* < 0.05, ***P* < 0.01, ****P* <0.001 by Kruskal-Wallis test with uncorrected Dunn’s test (n = 7 control vehicle, n = 5 control tasquinimod, n = 7 *Dnmt3a*^R878H/+^ vehicle, n = 5 *Dnmt3a*^R878H/+^ tasquinimod).
